# Morphological Changes and Mechanistic Analysis in Osteoblasts Induced by Long-Term Exposure to Radioactive Strontium

**DOI:** 10.64898/2026.11.15.766836

**Authors:** Zhen Wei, Shuo An, Jin Gao, Zhuojun Wu, Gengsheng Shi, Yiyao Cao, Hong Ren, Fengmei Cui, Liubin Li, Qiu Chen

## Abstract

Radioactive strontium-89 (^89^Sr) is preferentially deposited in bone tissue, where it directly interacts with osteoblasts; however, the morphological and molecular consequences of sustained ^89^Sr exposure remain poorly characterized. In this study, mouse pre-osteoblast MC3T3-E1 cells were induced to differentiate into osteoblasts and continuously exposed to 3.7 × 10³ Bq/mL ^89^Sr for nine consecutive passages. Morphological alterations were monitored throughout the exposure period, and the cells were subjected to RNA transcriptomic sequencing, non-targeted GC/MS-based metabolomics, and non-targeted LC/MS-based lipidomics, followed by integrative bioinformatics analysis. Prolonged ^89^Srr exposure markedly enlarged the cell nuclei and increased the number of intracellular vesicles in osteoblasts. Multi-omics co-enrichment analysis revealed metabolic perturbations centered on the “choline metabolism in cancer” and “fat digestion and absorption” pathways, while upregulated genes—including Hif1a, Egfr, Pdgfb, Egf, and Pdgfd—constituted key interaction modules underlying these alterations. Collectively, these findings suggest that ^89^Sr toxicity may be mediated through co-enriched pathways centered on choline metabolism and lipid digestion and absorption, driven by the upregulation of hub interaction modules comprising Hif1a, Egfr, Pdgfb, Egf, and Pdgfd. This study provides novel theoretical evidence for the toxicological mechanisms of radioactive strontium.

## 1 Introduction

Strontium is an alkaline earth metal with 21 isotopes ranging from ⁷⁸Sr to ⁹⁸Sr. Except for the naturally occurring stable isotopes ⁸⁴Sr, ⁹⁶Sr, ⁸⁷Sr, and ⁸⁸Sr, all others are radioactive isotopes. Among these, ⁸⁹Sr and ⁹⁰Sr are the most significant radioactive nuclides of strontium. Radioactive strontium-90 is a fission product of uranium and plutonium with a half-life of 28.8 years. It poses significant hazards to living organisms and ranks among the most persistent and dangerous radionuclides in the environment following nuclear accidents or nuclear weapon detonations^[1, 2]^. ⁸ ⁹ Sr is an artificial radionuclide that is often produced and used as a radiopharmaceutical. ⁸⁹Sr is a pure beta emitter with a maximum energy of 1.463 MeV and a half-life of 50.5 days^[3]^.Sr chloride is a widely used radiopharmaceutical worldwide for the treatment of severe pain caused by multiple osteoblastic bone metastases^[4]^, The principle is that due to its calcium-like chemical properties, intravenously administered Sr selectively accumulates in areas of active bone metabolism (particularly bone metastases), emitting beta particles to deliver internal radiation to the lesions. This effectively alleviates pain while minimizing damage to surrounding soft tissues and bone marrow due to its short range^[5]^.

After entering the skeleton, ⁸⁹Sr primarily deposits in the bone salt portion of the bone matrix, specifically the inorganic matrix. This deposition process involves two mechanisms: ion exchange and physiological osteogenesis. Over time, radioactive strontium participates in bone salt formation, thereby entering the crystals of bone inorganic salts. Radioactive strontium deposited within bones during physiological bone formation processes is difficult to remove. Long-term retention of radioactive strontium may cause damage to bone marrow hematopoietic tissue and abnormal bone formation, and may even induce bone cancer, leukemia, and associated peripheral soft tissue cancers^[6]^.

Previous studies have observed that ⁸⁹Sr induces hematologic toxicity in bone marrow, such as causing transient decreases in white blood cells and platelets^[4]^. It remains unclear whether long-term retention of ⁸⁹Sr in bone tissue affects osteoblast activity, thereby disrupting normal bone remodeling equilibrium. Detailed research is needed to investigate the skeletal toxicity effects and mechanisms of radioactive strontium, identifying key target molecules involved in its toxic mechanisms.This study observed morphological alterations in osteoblasts following prolonged exposure to radioactive strontium-89 and analyzed potential mechanisms through metabolic changes, identifying key molecules. The findings provide crucial theoretical data on the toxic effects and mechanisms of strontium accumulation in bone tissue after long-term retention.

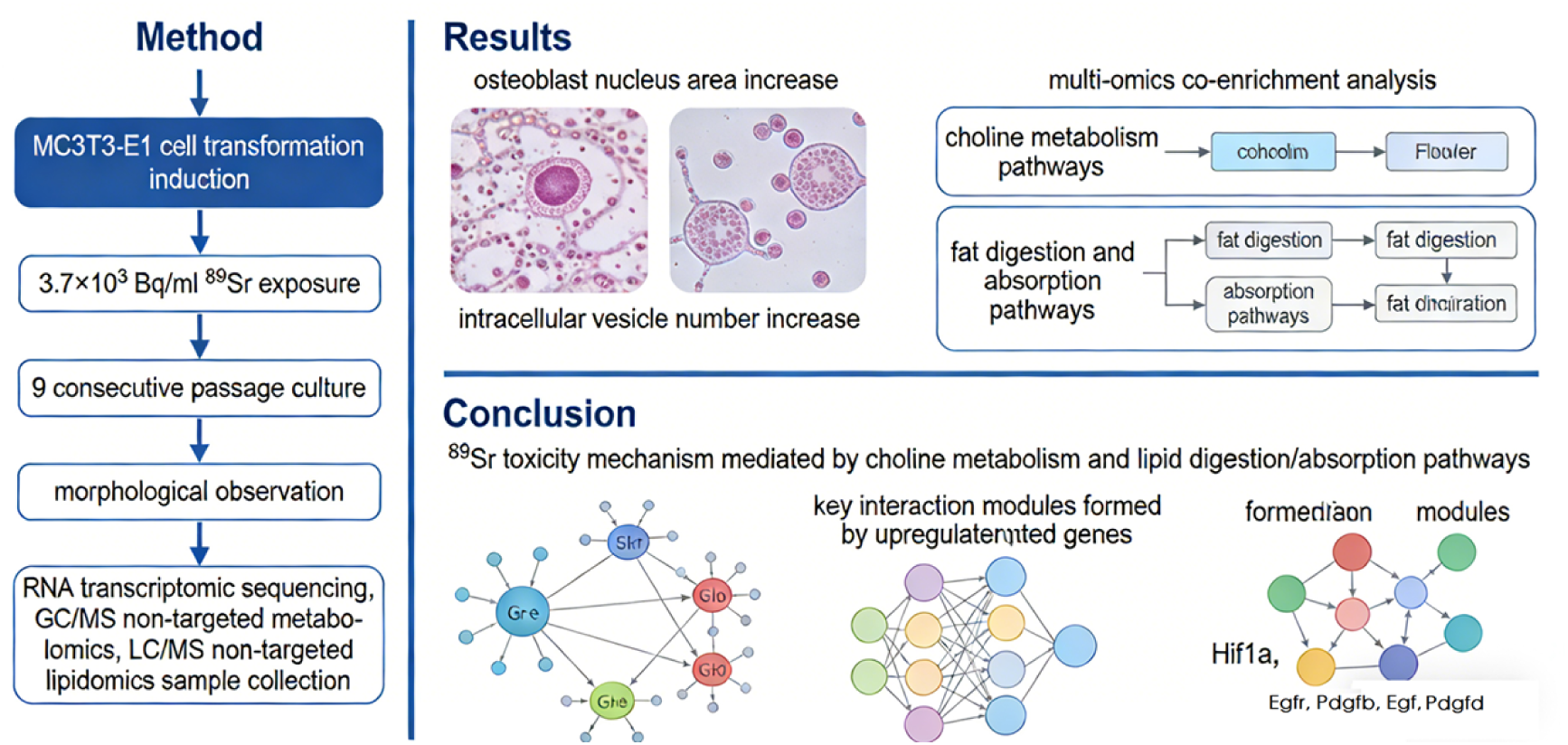

## 2 Materials and Methods

### 2.1 Cell culture

Mouse precursor osteoblasts MC3T3-E1 were induced to differentiate into osteoblasts, then exposed to 3.7 × 10³ Bq/ml radioactive strontium-89 and passaged continuously for 9 generations.Unexposed cells served as the control group (designated as NC). Cells exposed for the 6th generation (designated as P6) and 9th generation (designated as P9) were collected for subsequent experiments to establish a model of long-term strontium retention in osteoblasts. Collect cell samples from groups NC, P6, and P9 for testing of various indicators.

### 2.2 Cellular Morphology Observatio

Observe the cellular morphology of cells in the NC, P6, and P9 groups under an electron microscope, and use ImageJ software to calculate the nuclear area and the number of nuclear vesicles.

### 2.3 Transcriptome Sequencing Sample Preparation

Collect cell samples from the NC group and P9 group, with three biological replicates per group, and perform transcriptome sequencing. Samples were analyzed using Nanodrop for A260/A280 and A260/A230 ratios, combined with Agilent 2100 measurement of RIN values to verify RNA purity and integrity. Subsequently, library construction was performed by randomly shearing purified RNA into short fragments of 150–300 bp. Double-stranded cDNA was synthesized via reverse transcription, followed by PCR amplification to enrich the library. After passing quality control, high-throughput sequencing was completed on the Illumina platform. Filtering raw data yields clean reads. When a reference genome is available, sequence alignment is performed; when no reference genome exists, de novo assembly is conducted. Finally, gene expression levels are quantified using HTSeq, differentially expressed genes are identified via DESeq2, and GO/KEGG functional enrichment and PPI network analysis are performed to uncover biological significance.

### 2.4 GC/MS Non-Targeted Metabolomics Analysis

Collect cell samples from the NC and P9 groups for GC/MS non-targeted metabolomics analysis, with six biological replicates per group. After culturing cells to 85% confluence, remove the medium, digest with trypsin, and resuspend cells in an appropriate volume of pre-chilled 80% methanol aqueous solution. Incubate at –80 °C for one hour to fully precipitate proteins and extract metabolites. After centrifugation, collect the supernatant, remove the organic solvent by nitrogen blowdown concentration, and add methoxyamine hydrochloride-trimethylsilyl trifluoroacetamide for derivatization to enhance metabolite volatility. Derivatized samples were injected into a gas chromatography-mass spectrometry (GC-MS) system. Metabolites were separated using a capillary column and ionized via electron impact mode. Full-scan acquisition was performed to record metabolite retention times and characteristic ion signals. Raw data underwent peak identification, alignment, and quantitative peak area analysis using XCMS software, with baseline noise and batch effects removed. Metabolite qualitative annotation was performed using the NIST/Fiehn database. Differentially expressed metabolites were finally identified through multivariate statistical analyses including PCA and PLS-DA.

### 2.5 LC/MS Non-Targeted Lipidomics Analysis

Collect cell samples from the NC and P9 groups for LC/MS non-targeted lipidomics analysis, with six biological replicates per group. After culturing cells to 85% confluence, remove the medium, digest with trypsin, and resuspend cells in an appropriate volume of pre-chilled methyl tert-butyl ether-methanol extraction mixture. Vortex vigorously to thoroughly disrupt cells and extract lipids. After centrifugation, collect the supernatant organic phase. Concentrate by nitrogen blowdown, then redissolve in the initial mobile phase. Filter through a 0.22 μm organic-phase filter membrane. Samples were injected into the UPLC-MS system, where lipids were separated using a gradient elution with acetonitrile-isopropanol as the mobile phase. Both positive and negative ion modes were simultaneously acquired via the ESI source, combining full-scan and secondary fragmentation scanning to obtain lipid characteristic information. Raw data underwent peak processing and quantification using software such as Progenesis QI to eliminate noise and batch effects. Combined with qualitative annotation from the LipidMaps database, differential lipids were identified via PCA and OPLS-DA analysis, followed by metabolic pathway analysis.

### 2.6 Bioinformatics Analysis

Microarray gene expression profiles were extracted from the GEO database. Differential expression genes were analyzed using the online GEO2R tool with the following screening criteria: fold change ≥ 2.0 (log(FC) ≥ 1.0) and p-value < 0.05. Osteosarcoma data were selected from the TCGA database for analysis, and differential genes were screened for survival analysis.Using RStudio 4.5.2 with the ClusterGVis package, the mfuzz algorithm was applied to cluster different transcriptomic datasets. Kaplan-Meier curves were then plotted to visually represent trends, and the corrplot package was used to generate correlation plots between genes and metabolites. Cytoscape software was employed to visualize interaction networks among differentially expressed genes and metabolites.

### 2.7 Statistical Analysis

Statistical analysis of data was performed using GraphPad Prism 9.5 software. Quantitative data following a normal distribution are expressed as x ± s. Comparisons between two groups were performed using the t-test (assuming equal variances). P < 0.05 was considered statistically significant.

## 3 Result

### 3.1 Morphological Changes in Osteoblasts Following Strontium Toxicity

Cells from the NC, P6, and P9 groups were observed under a microscope. As shown in Figure 1, the results of cell morphology detection indicate that both the nuclear area and the number of nuclear vesicles in P9-generation cells significantly increased, suggesting that prolonged strontium exposure altered the morphology of osteoblasts.

**Figure 1.**
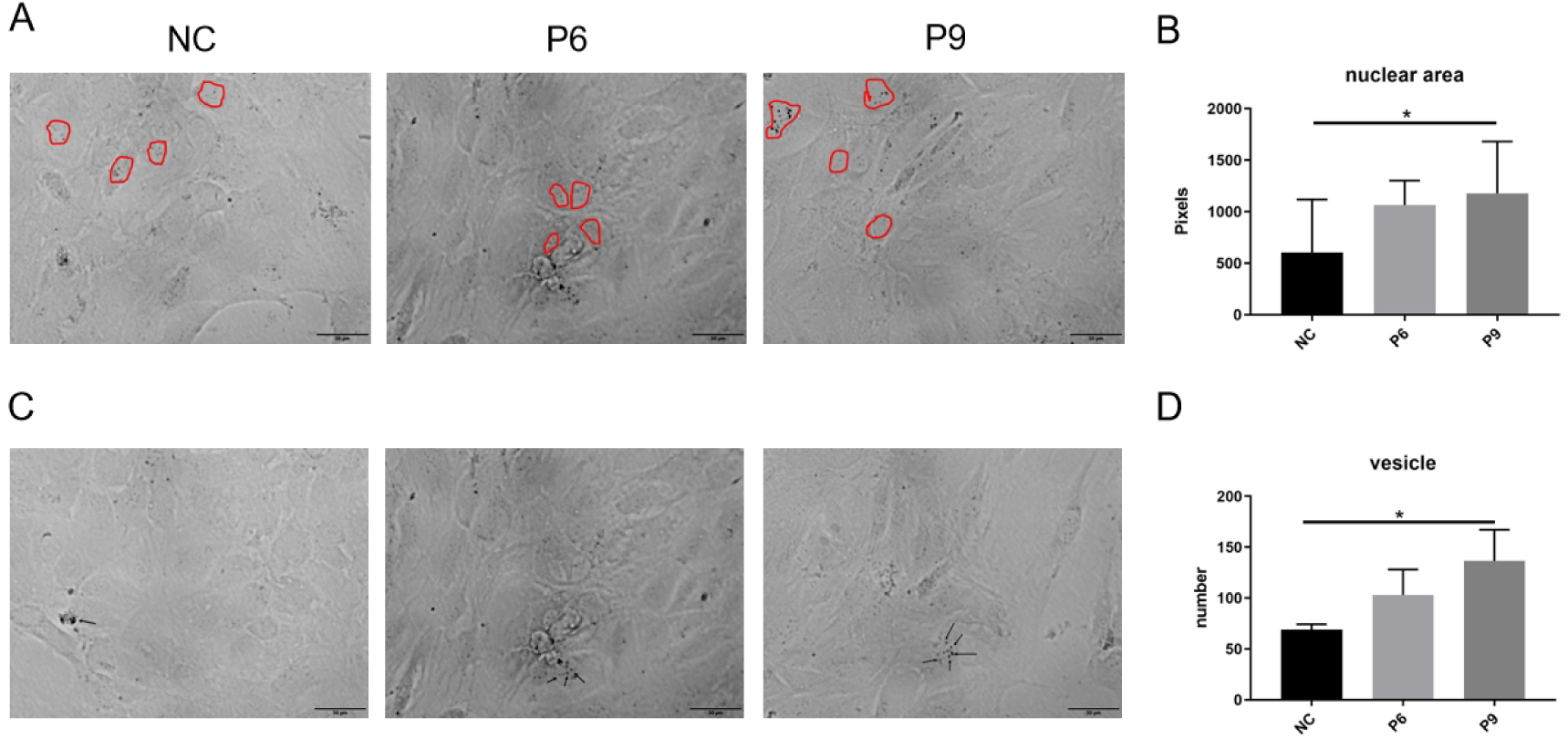
Morphological changes in osteoblasts following strontium staining. Panel A: Morphology of osteoblasts and nuclei. Panel B: Quantitative analysis of osteoblast nuclear area. Panel C: Intranuclear vesicles in osteoblasts. Panel D: Number of intranuclear vesicles.

### 3.2 Transcriptome Data Assembly Quality Analysis

We performed RNA-seq on the samples. Results showed good correlation among biological replicates, with R² ranging from 0.933 to 1.000; the correlation coefficient R² between control mice and irradiated mice ranged from 0.997 to 0.999 (Figure 2). The RNA-seq libraries from the 6 samples yielded clean read gene data ranging from 45.41 million to 48.93 million, with ValidBases (valid base mapping rate) between 86.78% and 88.54%. The Q30 base percentage for each sample ranged from 83.50% to 84.75% (Table S1). This indicates that the experimental sampling was reasonable and the RNA-seq data quality is reliable. Significant differences were observed between the two samples in both GC/MS non-targeted metabolomics analysis and LC/MS non-targeted lipidomics analysis, with the two coordinate points positioned relatively far apart on the score plot, confirming the reliability of sample selection.

**Figure 2.**
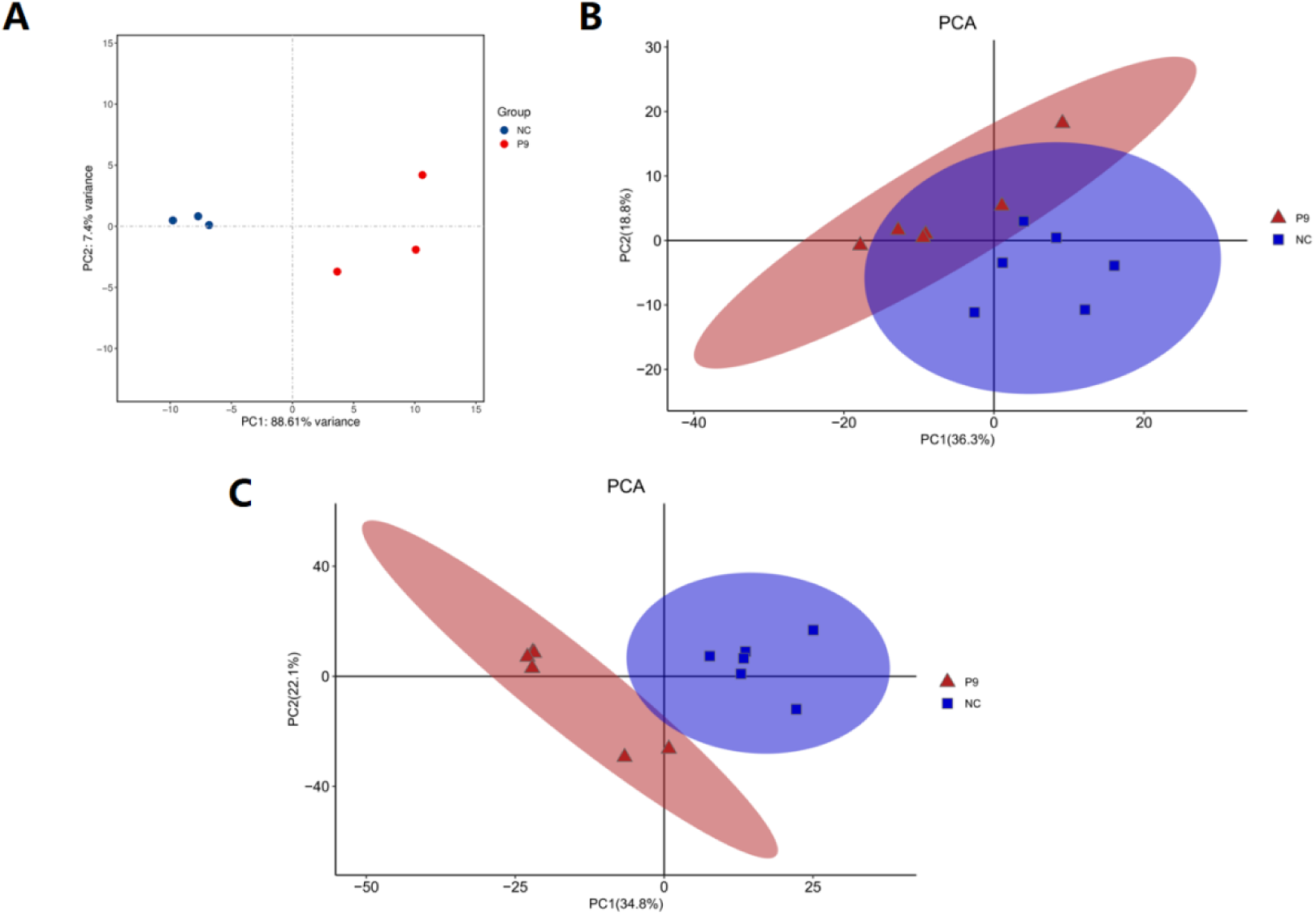
Principal Component Analysis (PCA) scoreplot of RNA-seq, GC/MS, and LC/MS data. Panel A. RNA-seq: Samples in the NC group exhibit significant intra-group clustering, while those in the PG group are completely separated from the NC group, indicating extremely significant differences in gene expression patterns between the two groups. Panel B: GC/MS non-targeted metabolomics. Samples in the NC group exhibit high intra-group clustering, while PG group samples partially overlap with the NC group but show overall trend separation, indicating distinct clustering patterns in metabolite composition between the two groups. Panel C: LC/MS non-targeted lipidomics. The NC group exhibits low intra-group dispersion, while PG group samples show clear separation from the NC group, indicating significant differences in lipid composition between the two groups.

### 3.3 Analysis of Differentially Expressed Results

All differential expression analyses were summarized based on the criteria of fold change > 2 and p < 0.05. Results revealed that differential gene analysis showed 357 genes were upregulated and 446 genes were downregulated; differential metabolite analysis indicated 4 metabolites were upregulated and 17 were downregulated; differential lipid metabolite analysis demonstrated 81 metabolites were upregulated and 44 were downregulated.

### 3.4 Enrichment analysis results for differentially expressed genes and metabolites

We performed GO analysis and KEGG analysis on differentially expressed genes. The results of the differential gene GO analysis revealed significant enrichment across multiple biological functions, including differential regulation at the molecular function, cellular component, and biological process levels. KEGG analysis results indicate that differentially expressed genes are primarily enriched in pathways such as the mTOR signaling pathway and the TCA cycle; Differential metabolite analysis revealed that the top 90 metabolites exhibited strong correlations, with absolute values of positive correlation coefficients ranging from 0.623 to 1.000. This indicates a pronounced tendency toward convergent changes among metabolites. KEGG pathway enrichment analysis revealed that differentially expressed metabolites primarily involved central carbon metabolism and amino acid metabolism pathways; Differential lipid metabolite analysis revealed that the top 90 metabolites exhibited strong positive (negative) correlations, with absolute correlation coefficients ranging from 0.623 to 1.000. This indicates significant clustering (divergence) among metabolites. KEGG pathway enrichment analysis revealed that differentially expressed metabolites primarily involved lipid-related pathways, such as glycerophospholipid metabolism and sphingolipid metabolism.

### 3.5 Relevance and Shared Pathway Analysis

To determine the correlation between differentially expressed metabolites and differentially expressed lipids with differentially expressed genes, a correlation analysis based on Pearson’s correlation coefficient was performed. A coefficient less than 0 indicates a negative correlation, while a coefficient greater than 0 indicates a positive correlation. Figure 5 The correlation analysis between the top 20 differentially expressed metabolites and lipids and the top 40 differentially expressed genes reveals that both groups exhibit positive and negative correlations with genes. However, the direction of these correlations (positive/negative) does not consistently align between the two groups. After performing KEGG enrichment analysis on differentially expressed metabolites, differentially expressed lipid metabolites, and differentially expressed genes separately, the shared enriched pathways between the two groups were identified. Among these, the top 20 metabolic pathways ranked by confidence for differentially expressed metabolites are: Protein digestion and absorption, arginine biosynthesis, basal cell carcinoma, aldosterone synthesis and secretion, PI3K-Akt signaling pathway, FoxO signaling pathway, cholesterol metabolism, morphine addiction, histidine metabolism, cGMP-PKG signaling pathway, choline metabolism in cancer, fat digestion and absorption, renin secretion, nitrogen metabolism, taurine and hyproline metabolism, arginine and proline metabolism, cancer-related pathways, glutathione metabolism, phenylalanine metabolism, biosynthesis of neomycin, kanamycin, and gentamicin. The top 20 metabolic pathways ranked by differential lipid metabolite reliability are: Choline metabolism in cancer, Glycerophospholipid metabolism, Retrograde endocannabinoid signaling, Insulin resistance, Neurotrophic factor signaling pathway, Herpesvirus associated with Kaposi’s sarcoma infection, AGE-RAGE signaling pathway in diabetic complications, leishmaniasis, adipokine signaling, fat digestion and absorption, amebiasis, regulation of lipolysis in adipocytes, thermogenesis, Th1 and Th2 cell differentiation, systemic lupus erythematosus, Egfr tyrosine kinase inhibitor resistance, ErbB signaling, natural killer cell-mediated cytotoxicity, and Th17 cell differentiation.

**Figure 3.**
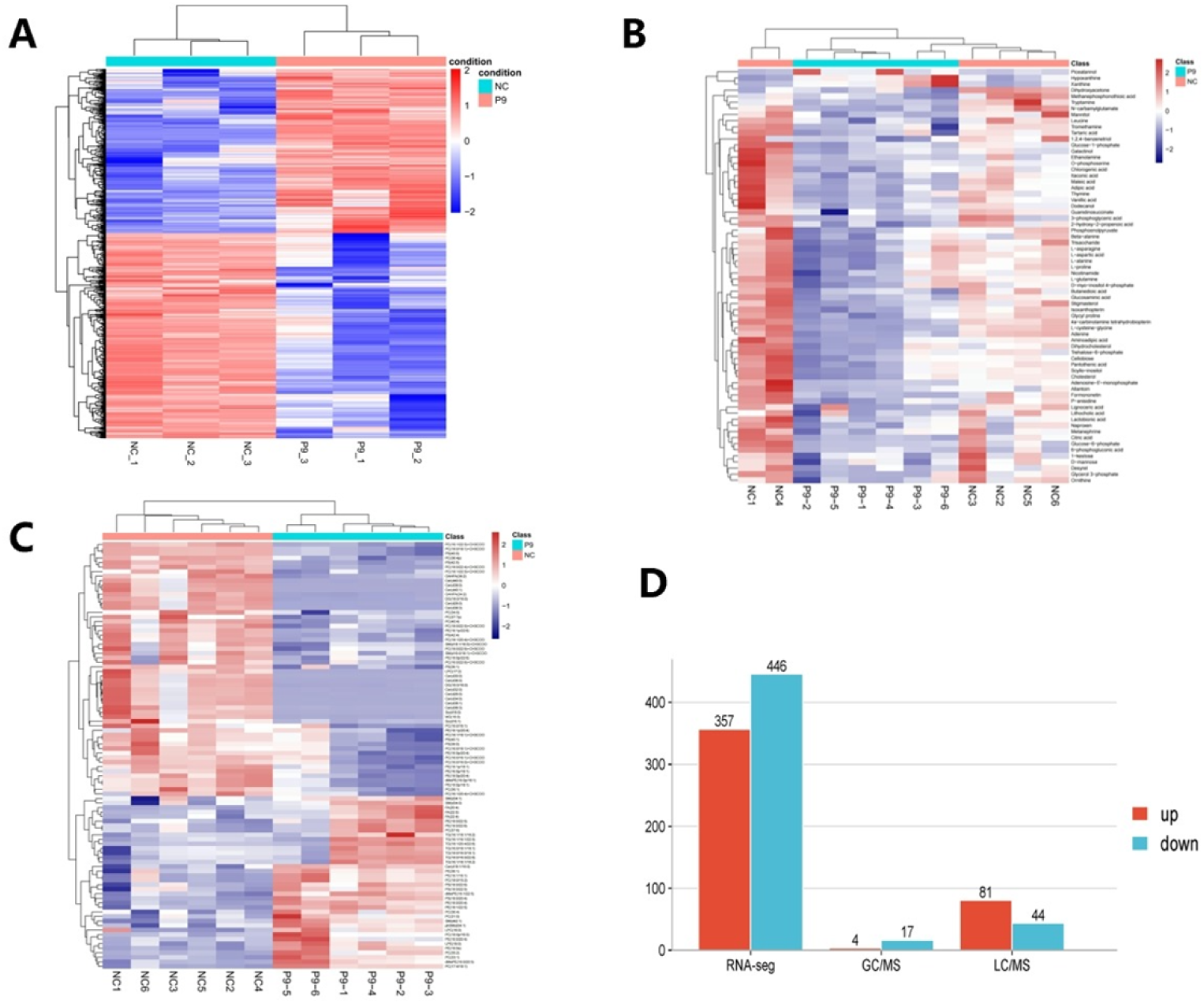
Analysis of differentially expressed results. Panel A: Differentially Expressed Genes (DEGs) Expression Heatmap. Colors are Z-score normalized, where red indicates genes with significantly higher relative expression levels than the sample average, blue indicates significantly lower levels, and white indicates levels close to the average. Panel B: GC-MS Metabolomics Differential Metabolite Expression Heatmap. Heatmap colors are Z-score normalized: red indicates metabolite relative expression significantly higher than the group’s average, blue indicates significantly lower than average, and white indicates expression close to average. Panel C: Differential lipid metabolite expression heatmap. Heatmap colors are Z-score normalized: red indicates lipid expression levels significantly higher than the group average, blue indicates levels significantly lower than average, and white indicates levels close to average. Panel D: Number of differentially regulated molecules in RNA-seq transcriptomics, GC/MS non-targeted metabolomics, and LC/MS non-targeted lipidomics: Red bars represent upregulated molecules, blue bars represent downregulated molecules.

**Figure 4.**
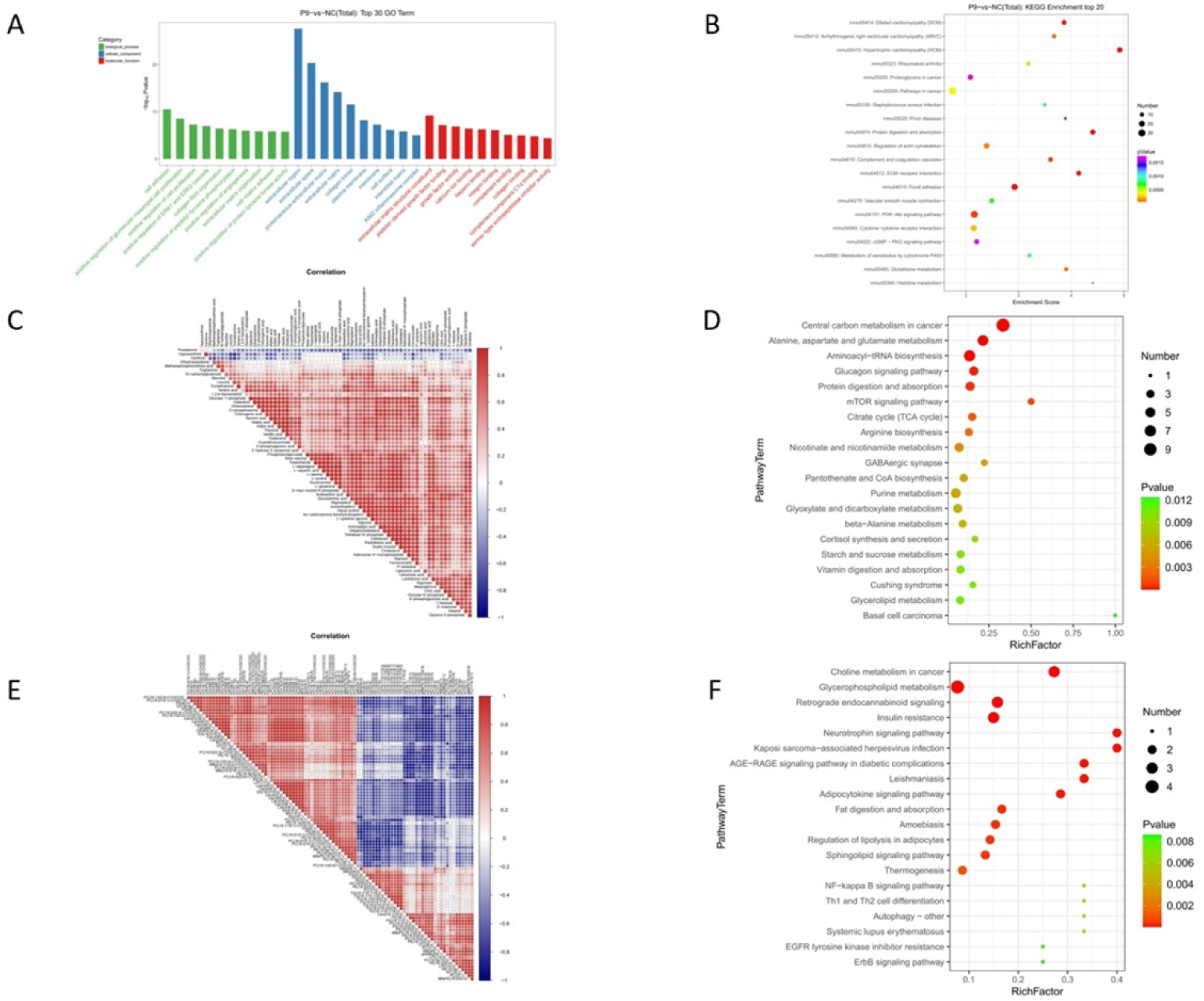
Enrichment analysis results for differentially expressed genes and metabolites. Panel A: Associative Clustering Diagram of Differentially Expressed Genes and Metabolites. This diagram illustrates the correlation distribution between differentially expressed genes and metabolites, where green represents biological process, blue represents cellular component, and red represents molecular function. The bar heights correspond to correlation coefficients, indicating the association trends between genes and metabolites. Panel B:KEGG enrichment bubble plot for differentially expressed genes. Bubble size corresponds to gene count, color indicates enrichment significance, and enrichment factors reflect the degree of enrichment for differentially expressed genes within pathways. Panel C: Correlation heatmap of differential metabolites. Colors in the figure correspond to correlation coefficients: red indicates positive correlation, blue indicates negative correlation. Panel D: KEGG pathway enrichment bubble plot for differentially expressed metabolites. Bubble size represents the number of differentially expressed metabolites within the pathway; bubble color corresponds to P-values, with red indicating P < 0.01 and green indicating P < 0.05. Darker colors denote higher significance. Panel E: Heatmap of correlations among differentially expressed lipid metabolites. Colors in the figure correspond to correlation coefficients: red indicates positive correlation, blue indicates negative correlation. Panel F: KEGG pathway enrichment bubble chart for differentially expressed lipid metabolites. Bubble size represents the number of differentially expressed lipids included in the pathway; bubble color corresponds to the P-value, with red indicating P < 0.01 and green indicating P < 0.05. Darker colors denote higher significance.

**Figure 5.**
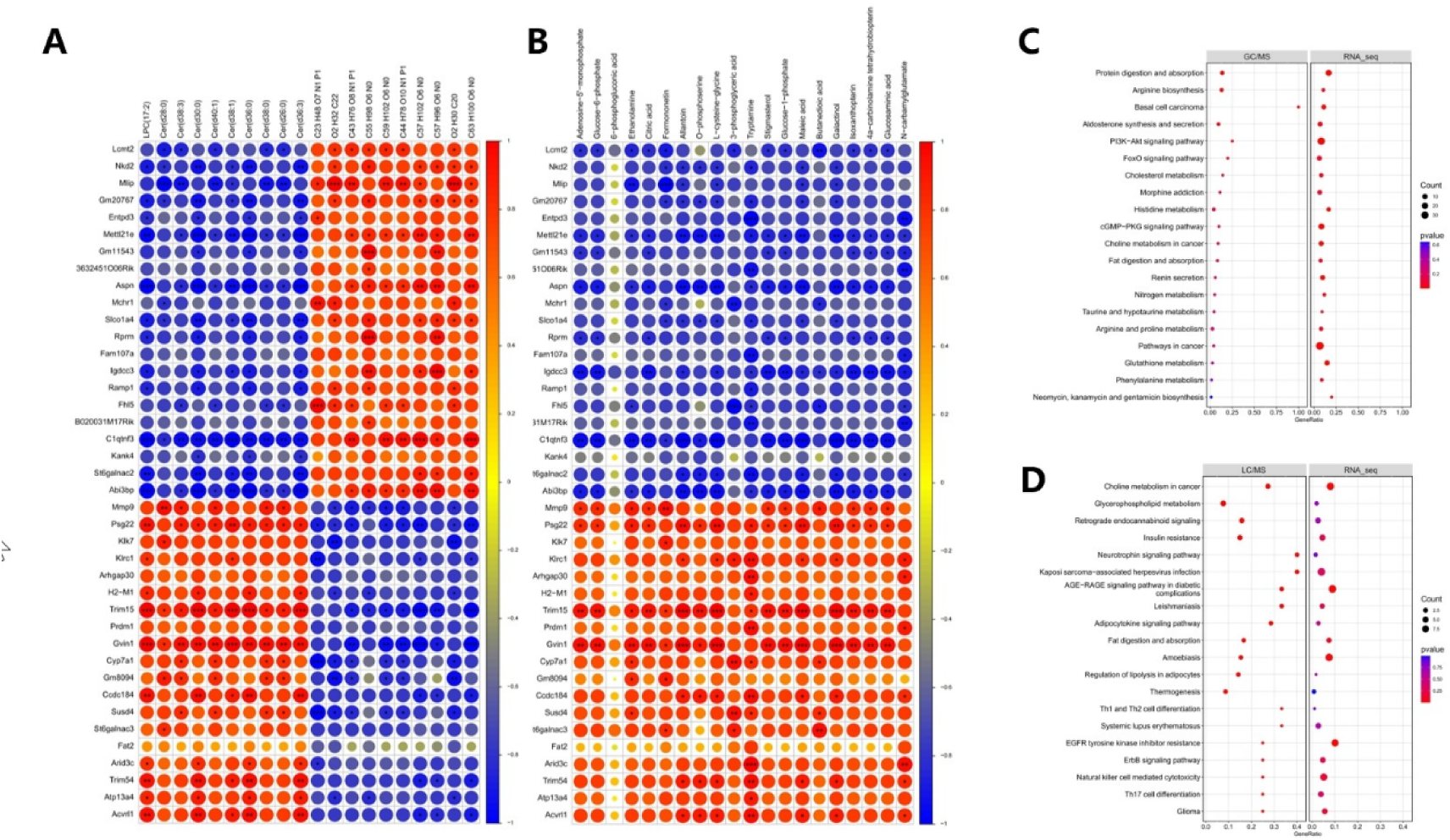
Relevance and Shared Pathway Analysis. Panel A: Heatmap of Pearson correlations between the Top 20 differential metabolites and the Top 40 differential genes. Dot color and size correspond to correlation coefficients, with red indicating positive correlation and blue indicating negative correlation. Results reveal both positive and negative correlations coexist between differential metabolites and differential genes. Panel B: Heatmap of Pearson correlations between the Top 20 differentially expressed lipid metabolites and the Top 40 differentially expressed genes. Dot color and size correspond to correlation coefficients, with red indicating positive correlation and blue indicating negative correlation. Results reveal both positive and negative correlations exist between differentially expressed lipid metabolites and genes. Moreover, the correlation directions between metabolites and genes are inconsistent. Panel C: GC-MS metabolomics differential analysis bubble plot. The horizontal axis represents enrichment factors, bubble size indicates gene count, and color corresponds to corrected P-values. The top 20 KEGG pathways for differential metabolites are displayed. Panel D: Bubble plot of enrichment pathways for differential lipid metabolites. The horizontal axis represents enrichment factors, bubble size indicates gene count, and color corresponds to corrected P-values. The top 20 KEGG pathways for differential lipid metabolites are displayed.

### 3.6 Correlation network of differentially expressed genes and differential metabolites in co-enriched pathways

Choline metabolism and fat digestion and absorption in cancer are pathways highly enriched for both differential metabolites and differential lipid metabolites, as well as differentially expressed genes. Therefore, we selected these pathways for further analysis. Correlation analysis revealed that the associations between metabolites and genes were predominantly positive correlations. Gene interaction analysis results indicate that the five genes Hif1a, Egfr, Pdgfb, Egf, and Pdgfd exhibit direct mutual interactions, and all five genes are enriched in the choline metabolism pathway in cancer. Among these, Hif1a serves as a core transcription factor in the hypoxic adaptation process, with its primary function being the regulation of target gene expression related to angiogenesis and energy metabolism. Egfr and its ligand Egf possess specific biological functions, capable of activating the PI3K-AKT pathway and the RAS-MAPK pathway. By activating these two pathways, they participate in biological processes such as cell proliferation, cell differentiation, and cellular damage repair. Pdgfb and Pdgfd both belong to the PDGF family. Their mechanism of action involves binding to the PDGFR β receptor to mediate vascular maturation and mesenchymal cell proliferation. Building upon this, we have mapped a pathway-gene-metabolite interaction network that visually illustrates the regulatory relationships among these three components, with results shown in Figure 6.

**Figure 6.**
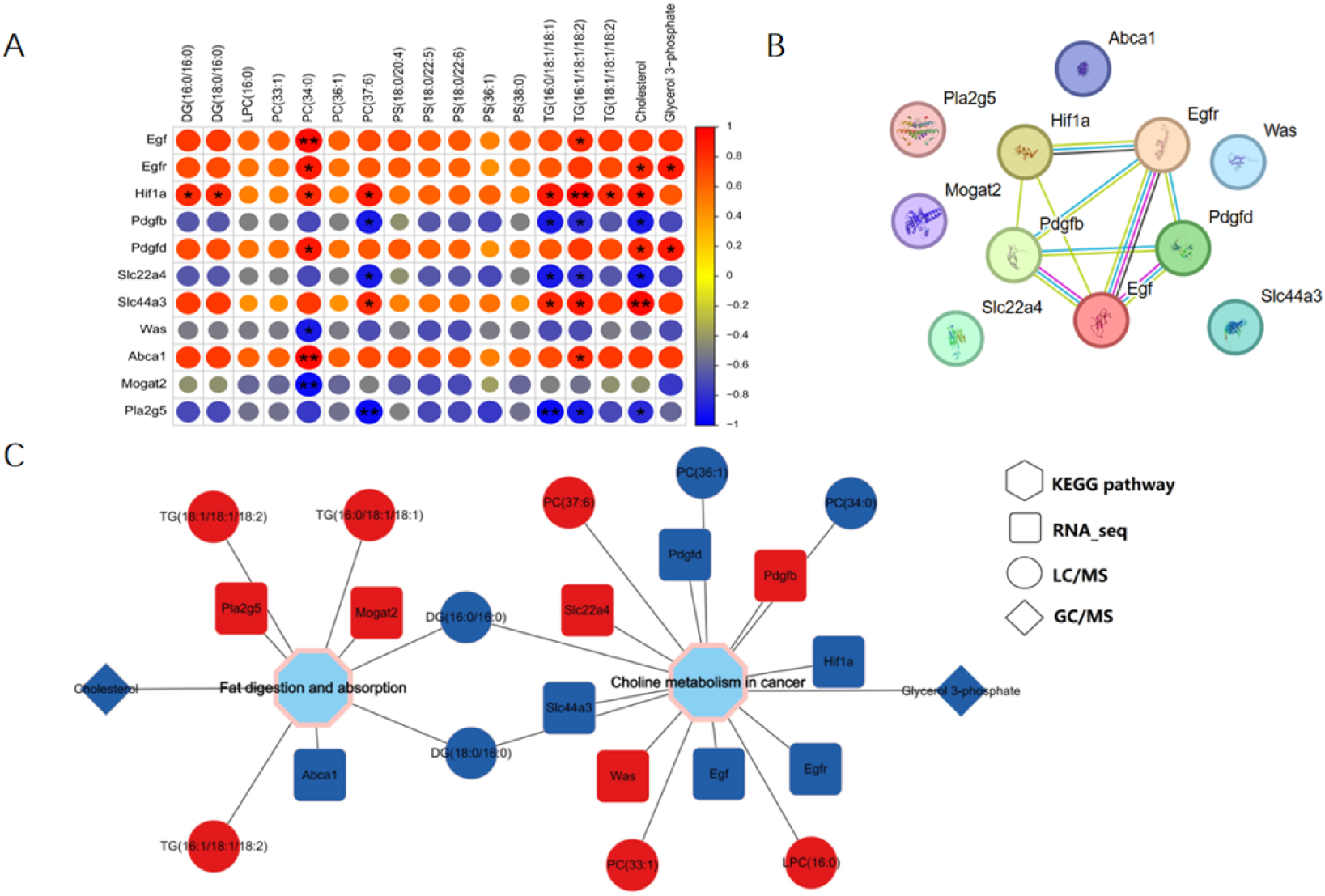
Panel A: Pearson correlation heatmap of core genes and metabolites: Dot color and size correspond to correlation coefficients, with red indicating positive correlation and blue indicating negative correlation. Results show both positive and negative correlations exist between core genes and metabolites. Panel B: Core gene interaction network. Nodes represent genes, and connections indicate interaction relationships. Panel C: Pathway-Gene-Metabolite Interaction Network. Central hexagons denote pathways. Square nodes correspond to RNA-seq molecules, circular nodes to LC-MS molecules, and diamond nodes to GC-MS molecules. Red nodes indicate upregulated molecules, while blue nodes indicate downregulated molecules.

**Figure 7.**
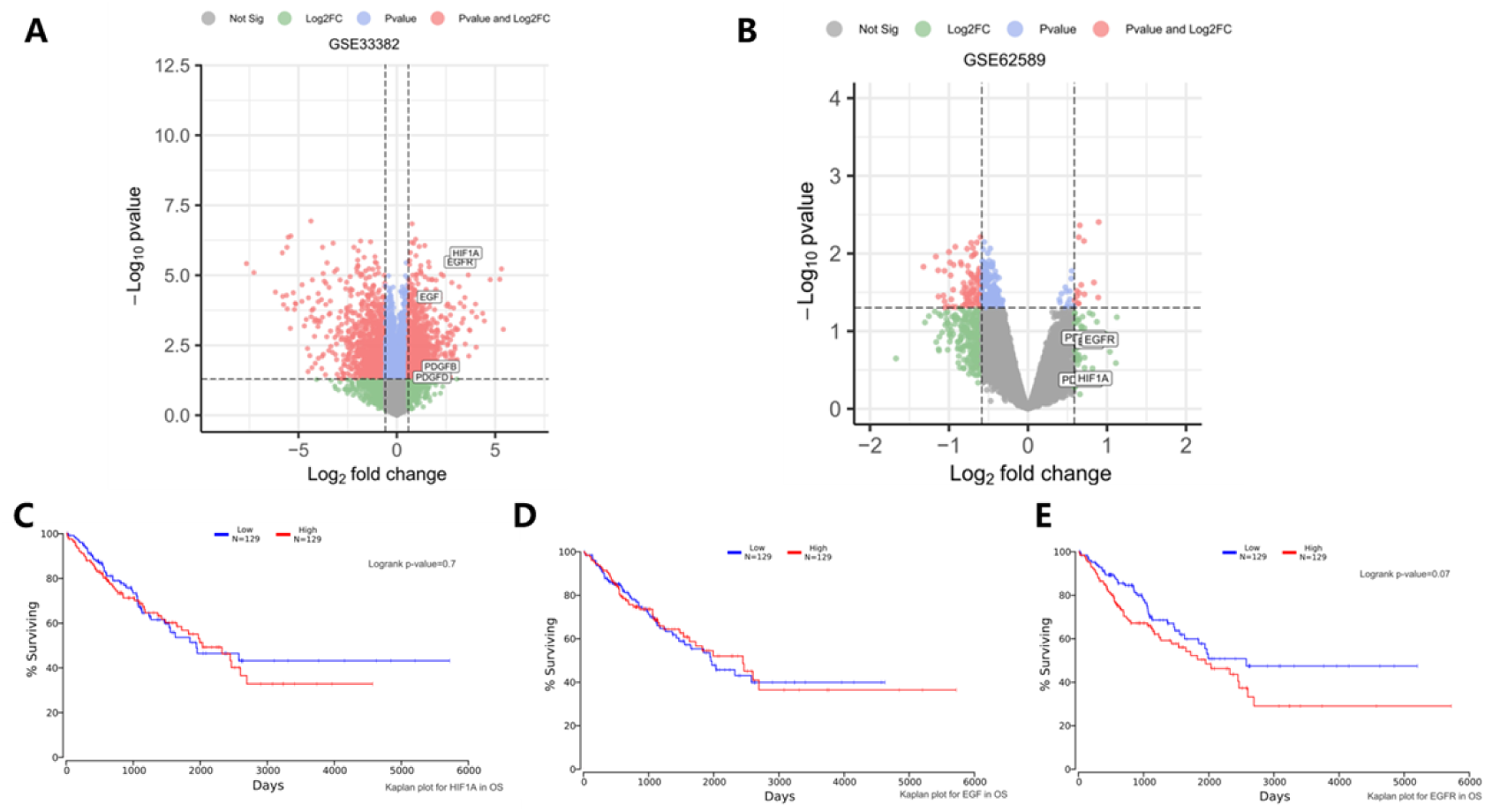
Panel A: Volcano plot of differentially expressed genes in the GSE33382 dataset (osteosarcoma). Different colors represent the screening results of differential analysis. The results show that genes such as EGF, HIF1A, and EGFR simultaneously meet the P-value and Log₂FC screening criteria in this dataset. Panel B: Volcano plot of differentially expressed genes in the GSE62589 dataset (osteoporosis): Different colors represent the screening results of differential analysis. The results show that genes such as EGF, HIF1A, and EGFR simultaneously meet both the P-value and Log₂FC screening criteria in this dataset. Panel C: Survival curves for HIF1A in the TCGA osteosarcoma dataset: Logrank test P-value = 0.7, indicating that HIF1A expression levels have no significant impact on survival duration in osteosarcoma patients. Panel D: Survival curves for EGF in the TCGA osteosarcoma dataset: Results indicate that EGF expression levels have no significant impact on survival duration in osteosarcoma patients. Panel E: Survival curves for EGFR in the TCGA osteosarcoma dataset: Logrank test P-value = 0.07, suggesting relatively longer median survival in the EGFR low-expression group.

### 3.7 Hif1a pathway participate in the development of osteoporosis or osteosarcoma

We searched the OMIM and GeneCards databases for target genes associated with osteoporosis or osteosarcoma, using a relevance score greater than 1 as the screening criterion. Ultimately, we identified genes related to both diseases, including HIF1A, EGF and EGFR. Detailed data is provided in Appendix 2. Subsequently, we identified osteosarcoma (GSE33382) and osteoporosis (GSE56815) datasets in the GEO database. Using R 4.5.2 software for differential gene analysis and applying a fold change threshold greater than 2, we found that all three genes were highly expressed in both datasets. Finally, to validate the roles of these three genes in osteosarcoma, we performed survival analysis on the osteosarcoma patient dataset from the TCGA database. The results showed that patients with low EGFR expression had a longer median survival compared to those with high expression, while the expression levels of the other two genes did not significantly affect survival in osteosarcoma.

## 4 Discussion

Radioactive strontium-89 is an artificial radionuclide and one of the most widely used radiopharmaceuticals globally for treating severe pain caused by multiple osteoblastic bone metastases. Despite its prominence and significance, research on the toxic effects and mechanisms of radioactive strontium remains severely lacking.

Cell morphology serves as a direct reflection of physiological function. This study revealed that after 89Sr-induced toxicity for nine generations (P9 group), osteoblasts exhibited significantly enlarged nuclear areas and increased numbers of nuclear vesicles. These morphological abnormalities suggest that cells may be in a state of stress or undergoing phenotypic transformation. Previously, research by Suh, K.S. et al. confirmed that nuclear morphology stability in osteoblasts is closely associated with bone matrix synthesis and mineralization functions. Nuclear structural abnormalities are typically accompanied by disrupted gene transcription regulation and abnormal expression of functional proteins^[7]^.

Transcriptome results indicate that ⁸⁹Sr exhibits long-term retention in osteoblasts, extensively disrupting their gene expression networks. Further analysis of altered metabolome and lipidome expression reveals that post-transcriptional regulation of gene expression, feedback loops in metabolite synthesis or degradation, and signaling functions of lipid molecules may all be implicated^[8]^.

Based on Pearson correlation coefficient analysis, differentially expressed metabolites primarily clustered in pathways such as protein digestion and absorption, arginine biosynthesis, and the PI3K-Akt signaling pathway. Differentially expressed lipid molecules, meanwhile, concentrated in pathways including glycerophospholipid metabolism, retrograde endocannabinoid signaling, and insulin resistance. These pathways respectively involve multiple core physiological processes such as nutrient metabolism, cellular signaling, and energy metabolism regulation. The PI3K-Akt signaling pathway plays a crucial role in the aforementioned physiological processes. Current research has demonstrated that this pathway serves as a key regulator of osteoblast proliferation, differentiation, and survival. Dysfunction of this pathway can directly inhibit osteogenic differentiation and promote apoptosis ^[9–11]^. Additionally, disruptions in lipid metabolism pathways, such as abnormalities in glycerophospholipid metabolism, may further exacerbate functional impairment in osteoblasts by affecting processes including cell membrane structural stability and the release of signaling molecules^[12]^.

Multi-omics co-enrichment analysis is an effective method for identifying core regulatory pathways. Our study reveals that choline metabolism and fat digestion and absorption in cancer are key pathways co-enriched by differentially expressed genes, metabolites, and lipid molecules, indicating that these pathways play a crucial role in osteoblast damage induced by ⁸⁹Sr radiation exposure.

Research has confirmed that disruptions in the lipid digestion and absorption pathway are associated with abnormal energy metabolism in osteoblasts. Fatty acids generated through lipid metabolism provide a crucial energy supply for osteoblasts, and these fatty acids also function as signaling molecules to regulate osteoblast differentiation^[13]^.This indicates that when fat digestion and absorption are impaired, osteoblast energy supply decreases and bone matrix synthesis is reduced. Additionally, impaired fat digestion and absorption leads to lipid accumulation, triggering lipotoxic stress that damages osteoblasts. Concurrently, the significant overexpression of choline metabolism pathways in cancer represents another key finding of this study. Choline contributes to the composition of phospholipids in cell membranes, while its metabolites—such as acetylcholine and phosphatidylcholine—also function as signaling molecules regulating cellular proliferation, differentiation, and apoptosis^[14, 15]^.

Our analysis of gene interactions reveals that the five genes Hif1a, Egfr, Pdgfb, Egf, and Pdgfd, originating from the pathway “Choline metabolism in cancer,” all interact with each other and are differentially expressed. This suggests that these genes form a key gene module and may represent potential targets for ⁸⁹Sr radiation exposure. HIF-1 α (hypoxia-inducible factor 1α) in mice is a crucial transcription factor regulating hypoxic adaptation. It activates during hypoxia and initiates the expression of a series of downstream target genes, promoting angiogenesis and modulating cellular energy metabolism patterns to sustain cell survival. It plays a vital role in embryonic development, tumor microenvironment adaptation, and tissue ischemic repair. ^[16]^. Egfr (epidermal growth factor receptor) is a transmembrane tyrosine kinase receptor. Upon ligand binding, it activates downstream signaling pathways such as PI3K-AKT and RAS-MAPK, participating in the regulation of proliferation, differentiation, migration, and apoptosis in mouse cells. Egfr is essential for mouse epidermal renewal, hair follicle development, and lung tissue maturation, and is also closely implicated in tumorigenesis and tumor progression^[17, 18]^. Pdgfb (Platelet-derived growth factor B) belongs to the PDGF family and is primarily secreted by platelets and vascular endothelial cells. It binds to the PDGFRβ receptor and mediates the recruitment and proliferation of vascular smooth muscle cells (VSMCs) and pre-capillary plasma cells (PCs). While playing a crucial role in vascular maturation and stability, it also promotes the proliferation of mesenchymal cells during embryonic organogenesis and tissue repair following injury^[19, 20]^. Egf (Epidermal growth factor) is one of the key ligands for Egfr. By specifically binding to Egfr and activating receptor signaling, it not only potently promotes epithelial cell proliferation and migration but also participates in tissue repair following injury, playing a crucial role in maintaining epithelial tissue integrity^[21]^. Pdgfd (Platelet-derived growth factor D) is also a member of the PDGF family, with PDGFRβ as its receptor. While it shares functional overlap with Pdgfb, their roles differ in emphasis. Pdgfd primarily functions during late embryonic development and in adult tissues, regulating the activation and proliferation of mesenchymal cells. It participates in physiological and pathological processes such as organ fibrosis, vascular stability, and adipose tissue development ^[22, 23]^. The osteoblast toxicity induced by ⁸⁹Sr is highly likely mediated through the regulatory network established by the aforementioned five genes. Simultaneously, activation of Hif1a promotes cell proliferation to counteract adverse effects by upregulating the expression of genes such as Egfr and Pdgfb, as well as through the PI3K-AKT pathway. However, prolonged abnormal activation may further induce phenotypic changes in osteoblasts and cellular dysfunction, ultimately impairing bone formation or leading to malignant transformation. It can be inferred that the cellular morphological and functional abnormalities observed in this study are caused by pathological factors, and this finding opens new avenues for future research into the mechanisms underlying the long-term toxic effects of radionuclides on bone tissue.

Our findings elucidate the molecular mechanisms underlying the toxic effects of radioactive strontium from a metabolic perspective, identifying key genes and metabolites involved. This research provides candidate intervention targets for biological protection against the toxic effects of radioactive strontium.

## Author contributions statement

Conceptualization, Qiu Chen and Liubin Li; methodology, Zhen Wei, Jin Gao and Zuojun Wu; investigation, Zhen Wei, Jin Gao, Zuojun Wu, Gengsheng Shi, Yiyao Cao and Hong Ren; formal analysis, Gengsheng Shi and Fengmei Cui; data curation, Fengmei Cui; writing— original draft preparation, Zhen Wei, Jin Gao and Zuojun Wu; writing—review and editing, Qiu Chen and Liubin Li; visualization, Gengsheng Shi and Fengmei Cui; supervision, Qiu Chen and Liubin Li; funding acquisition, Qiu Chen and Liubin Li; All authors have read and agreed to the published version of the manuscript.

## Acknowledgements

The author(s) disclosed receipt of the following financial support for the research, authorship, and/or publication of this article: This work was supported by the Natural Science Research Projects of Colleges and Universities in Jiangsu Province (grant numbers 20KJB310007), the Priority Academic Program Development of Jiangsu Higher Education Institutions (PAPD), the Suzhou Key Disciplines (No. SZXK202506), the Collaborative Innovation Center of Radiological Medicine of Jiangsu Higher Education Institutions.

## Supplements

**Table 1.**
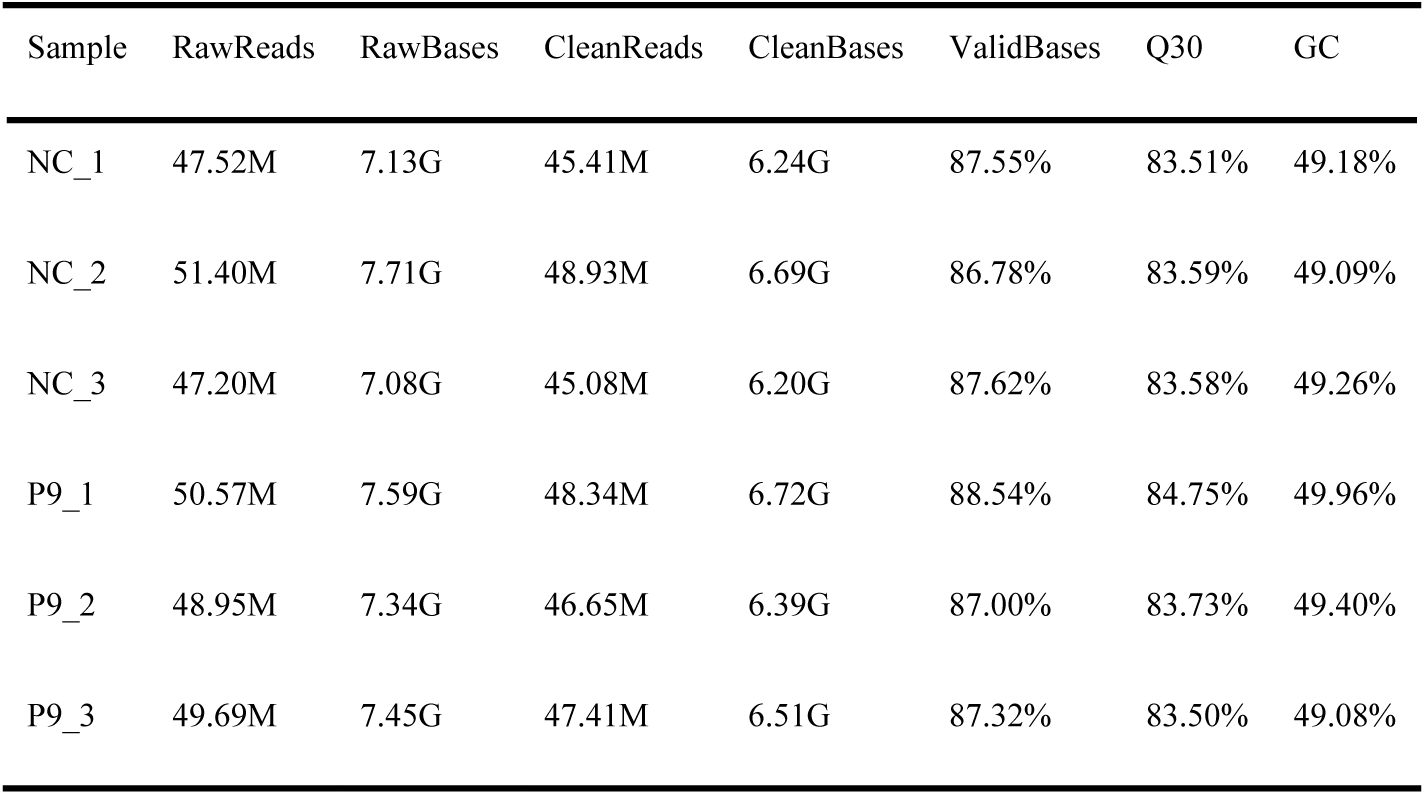
Sequencing Data Quality Results Table.

**Table 2.** Summary of Common Target Genes for Osteoporosis and Osteosarcoma.

| **Gene Symbol** | **Description** | **Uniprot ID** | **Gifts** | **GC Id** | **Relevance score** |
|---|---|---|---|---|---|
| **osteosarcoma** |  |  |  |  |  |
| HIF1A | Hypoxia Inducible Factor 1 Subunit Alpha | Q16665 | 62 | GC14P061695 | 14.19 |
| EGFR | Epidermal Growth Factor Receptor | P00533 | 67 | GC07P055019 | 9.08 |
| PDGFB | Platelet Derived Growth Factor Subunit B | P01127 | 63 | GC22M091586 | 6.64 |
| EGF | Epidermal Growth Factor | P01133 | 64 | GC04P109912 | 5.19 |
| PDGFD | Platelet Derived Growth Factor D | Q9GZP0 | 55 | GC11M103907 | 1.59 |
| **Osteoporosis** |  |  |  |  |  |
| EGF | Epidermal Growth Factor | P01133 | 64 | GC04P109912 | 5.98 |
| HIF1A | Hypoxia Inducible Factor 1 Subunit Alpha | Q16665 | 62 | GC14P061695 | 3.18 |
| EGFR | Epidermal Growth Factor Receptor | P00533 | 67 | GC07P055019 | 1.94 |
| PDGFB | Platelet Derived Growth Factor Subunit B | P01127 | 63 | GC22M091586 | 0.28 |

